# *C. elegans* paraoxonase-like proteins control the functional expression of DEG/ENaC mechanosensory proteins

**DOI:** 10.1101/040337

**Authors:** Yushu Chen, Shashank Bharill, Zeynep Altun, Robert O’Hagan, Brian Coblitz, Ehud Y. Isacoff, Martin Chalfie

**Author notes:** To whom correspondence should be sent at: Department of Biological Sciences, 1012 Fairchild, MC#2446, Columbia University, 1212 Amsterdam Avenue, New York, NY 10027.

## Abstract

*Caenorhabditis elegans* senses gentle touch via a mechanotransduction channel formed from the DEG/ENaC proteins MEC-4 and MEC-10. An additional protein, the paraoxonase-like protein MEC-6, is essential for transduction, and previous work suggested that MEC-6 was part of the transduction complex. We found that MEC-6 and a similar protein, POML-1, reside primarily in the endoplasmic reticulum and do not colocalize with MEC-4 on the plasma membrane in vivo. As with MEC-6, POML-1 is needed for touch sensitivity, for the neurodegeneration caused by the *mec-4(d)* mutation, and for the expression and distribution of MEC-4 in vivo. Both proteins are likely needed for the proper folding or assembly of MEC-4 channels in vivo as measured by FRET. MEC-6 detectably increases the rate of MEC-4 accumulation on the *Xenopus* oocyte plasma membrane. These results suggest that MEC-6 and POML-1 interact with MEC-4 to facilitate expression and localization of MEC-4 on the cell surface. Thus, MEC-6 and POML-1 act more like chaperones for MEC-4 than channel components.

**Abbreviations used**
TRNs: touch receptor neurons
PONs: paraoxonases
POML-1: <u>p</u>ara<u>o</u>xonase
<u>M</u>EC-6-<u>l</u>ike: gene 1
ER: endoplasmic reticulum

## INTRODUCTION

Gentle touch is sensed in the nematode *Caenorhabditis elegans* by six touch receptor neurons (TRNs) (Chalfie and Sulston, 1981). Touch is transduced in the TRNs by the activation of a trimeric channel formed by two DEG/ENaC (degenerin/epithelial sodium channel) proteins MEC-4 and MEC-10 (O’Hagan *et al*., 2005; Arnadóttir *et al*., 2011; Chen *et al*., 2015). Previous work from our lab suggested that another protein, MEC-6, was also part of the mechanosensory channel complex in the TRNs, since it colocalized with MEC-4 in TRN neurites and co-immunoprecipitated with it in heterologous cells (Chelur *et al*., 2002).

The sequence of MEC-6 and several other predicted *C. elegans* proteins (Chelur *et al*., 2002) is similar to that of the three mammalian paraoxonases (PON1-PON3). Human PON1 and PON3 are serum proteins that contribute to HDL particles (Mackness and Walker, 1988; Reddy *et al*., 2001). In contrast, human PON2, which is ubiquitously expressed (Mochizuki *et al*., 1998), localizes to the endoplasmic reticulum (Horke *et al*., 2007; Rothem *et al*., 2007). The exact function of these proteins is unclear, but they prevent lipid peroxidation (Aviram *et al*., 1998; Shih *et al*., 1998; Ng *et al*., 2001; Reddy *et al*., 2001; Besler *et al*., 2011; Devarajan *et al*., 2011; Huang *et al*., 2013) and PON1, but not PON2 or PON3, degrades the organophosphate paraoxon (Smolen *et al*., 1991; Davies *et al*., 1996; Stevens *et al*., 2008).

The localization and function of the mammalian paraoxonases suggest that MEC-6 may not be an integral component of the mechanosensory transduction complex, but interacts with the channel-forming subunits elsewhere in the cell. Indeed, we recently found that MEC-6 does not colocalize with MEC-4 either on the plasma membrane of *Xenopus* oocytes or in TRN neurites (Chen *et al*., 2015). These results when coupled to studies in *Drosophila melanogaster*, which has mechanosensory DEG/ENaC proteins (Liu *et al*., 2003; Zhong *et al*., 2010; Gorczyca *et al*., 2014; Guo *et al*., 2014; Mauthner *et al*., 2014), but no obvious MEC-6-like proteins (Hicks *et al*., 2011), led us to reinvestigate the role of MEC-6 in *C. elegans* touch sensitivity.

Here we show that MEC-6 and a second paraoxonase-like protein that is expressed in the TRNs, POML-1 [<u>p</u>ara<u>o</u>xonase and <u>M</u>EC-6-<u>l</u>ike gene 1, previously named K11E4.3 (Topalidou and Chalfie, 2011)], primarily reside in the endoplasmic reticulum (ER) of the TRN cell body. Our results suggest that MEC-6 and POML-1 are important for MEC-4 production and localization.

## RESULTS

### MEC-6 and POML-1 localize to the TRN endoplasmic reticulum

The POML-1 sequence is 30% identical and 42% similar to that of MEC-6. Both MEC-6 (Chelur *et al*., 2002) and POML-1 contain a transmembrane domain and a nematode-specific region of 15 amino acids in the N-terminus (Figure S1A). In contrast to MEC-6, which is expressed in many cells (Chelur *et al*., 2002), POML-1 was expressed in only a few neurons. In addition to the six TRNs (Topalidou and Chalfie, 2011), POML-1 was found in the IL1, AIM, ALN, and BDU neurons (Figure S1B).

Rescuing and tagged translational fusions of MEC-6 and POML-1 are primarily expressed in the TRN cell body, with weak diffuse expression and puncta in the proximal TRN neurite (Figure S1C, and Chen *et al*., 2015). Using antibodies against tagged proteins, we usually saw POML-1 further along the TRN neurites (up to 100 μm of the approximately 400 μm neurite) than MEC-6 (Figure S1D). In the cell body, translational fusions for both proteins formed a perinuclear mesh-like structure that colocalized with markers for the ER [YFP::TRAM-1 and YFP::PISY-1, which label rough ER and general ER, respectively (Rolls *et al*., 2002)] and each other (Figure 1, A-C), but not with a marker for the Golgi apparatus (AMAN-2::YFP; Figures 1D, Figures 1E, and S1E). [POML-1 partially overlapped with the Golgi marker in 3 of 10 cells (Figure S1E)]

**Figure 1.**
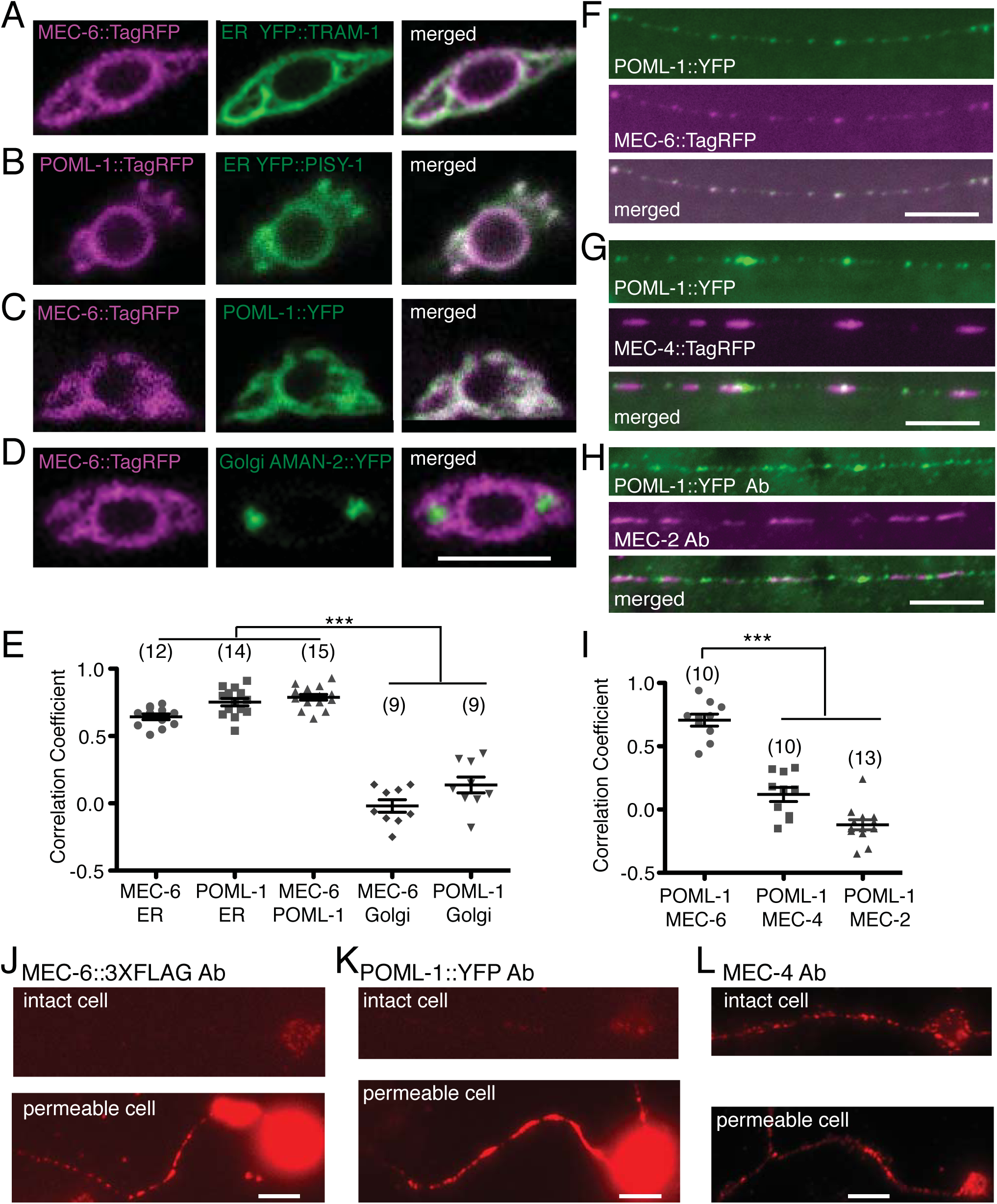
TRN expression of MEC-6 and POML-1. Confocal sections of TRN cell bodies of (A) MEC-6::TagRFP and the ER marker YFP::TRAM-1, (B) POML-1::TagRFP and the ER marker YFP::PISY-1, (C) MEC-6::TagRFP and POML-1::YFP, and (D) MEC-6::TagRFP and the Golgi marker AMAN-2::YFP, and their correlation coefficient (E). Scale bars = 5 μm (here and in F, G, H, J, K, and L). The number of examined TRNs is given in parenthesis (E and I). Symbolism for significance here and in all subsequent figures is described in the Materials and Methods. Neurite expression of (F) POML-1::YFP and MEC-6::TagRFP, (G) POML-1::YFP and MEC-4::TagRFP, and (H) POML-1::YFP and MEC-2 and their correlation coefficient (I). Anti-GFP and anti-MEC-2 antibodies (Ab) were used to label the proteins in (H). (J) MEC-6::3XFLAG expression as detected by an anti-FLAG antibody in intact (upper panel) and permeabilized (lower panel) cultured TRNs. (K) POML-1::YFP expression as detected by an anti-GFP antibody in intact (upper panel) and permeabilized (lower panel) TRNs in culture. The faint immunofluorescence in intact cells (J and K) was not specific because it was often observed in cells that did not express MEC-6::3XFLAG or POML-1::YFP. Images in J and K are representative of 40 cells examined in two independent experiments. (I) MEC-4 expression detected with an anti-MEC-4 antibody that recognizes the extracellular domain in intact (upper panel) and permeabilized (lower panel) cultured TRNs. Images are representative of 20 cells examined in two independent experiments.

When seen, MEC-6 and POML-1 puncta colocalized in the proximal TRN neurite (Figure 1F). As with MEC-6 (Chen *et al*., 2015), POML-1 puncta differed from and did not colocalize with MEC-4::TagRFP or MEC-2 puncta in TRN neurites (Figure 1, G-I) although some overlap was observed. [General ER but not rough ER is also present in TRN neurites (Figure S1F; Rolls *et al*., 2002). In general, POML-1 puncta (of the cells represented in Figure 1, F and G) were smaller and closer together than the MEC-4 and MEC-2 puncta: POML-1 puncta were 0.90 ± 0.02 μm wide and were separated by 1.17 ± 0.08 μm (n = 25 PLM neurites), and MEC-2 puncta were 1.95 ± 0.05 μm wide and were separated by 3.86 ± 0.13 μm [n = 30 PLM neurites; p<0.0001 for both puncta width and the distance between puncta for POML-1::YFP and MEC-2].

Consistent with the ER localization of MEC-6 and POML-1, we found that both proteins were absent from the TRN surface. Previous work suggested that MEC-6 was a membrane protein that extended its C-terminus extracellularly (Chelur *et al*., 2002). Specifically, LacZ fused to the C-terminus of MEC-6 produced no β-galactosidase activity unless a synthetic transmembrane domain was inserted between MEC-6 and LacZ (LacZ only produces β-galactosidase activity intracellularly). We found that POML-1 acted similarly (Figure S1G). This result suggests that the C-termini of MEC-6 and POML-1 are not located in the cytoplasm, but are either outside the cell or in the lumen of an internal organelle. MEC-6 was glycosylated when expressed in CHO cells, and C-terminally HA-tagged MEC-6 was detected by surface immunostaining against HA tags in CHO cells (Chelur *et al*., 2002). However, in cultured TRNs from wild type embryos (using antibodies directed against C-terminal tags of MEC-6 and POML-1), we were only able to detect the proteins when the cells were permeabilized, indicating the absence of both MEC-6 and POML-1 expression on the cell surface (Figure 1, J and K, n = 40 cells for each). On the contrary, an antibody recognizing the extracellular region of MEC-4 detected clear expression of MEC-4 on the cell surface in intact cells (Figure 1L, n = 20 cells). Since MEC-6 and POML-1 could be expressed, albeit weakly, on the surface of *Xenopus* oocytes (Figure S1H, and Chen *et al*., 2015) and CHO cells in previous experiments (Chelur *et al*., 2002), either control over the subcellular localization is tighter in the TRNs or a small, undetected amount of MEC-6 and POML-1 goes to the TRN surface. Most of MEC-6 and POML-1, however, was detected in the TRN ER.

The above fusion constructs, including *poml-1::yfp*, *poml-1::tagrfp*, *mec-4::tagrfp, mec-6::3Xflag*, and *mec-6::tagrfp* produced functional products since they could rescue *poml-1, mec-4*, and *mec-6* null mutations (Figure S1I, and Chen *et al*., 2015).

### POML-1 affects the function of MEC-4

The failure of POML-1 to colocalize with MEC-4 suggests that POML-1 may not function directly in transduction. To study the role of *poml-1* gene, we first generated *poml-1* null alleles. Using Mos1-mediated gene deletion (Frokjaer-Jensen *et al*., 2010; Frokjaer-Jensen *et al*., 2012), we obtained two null mutations, *u881* and *u882*, which lacked the entire *poml-1* coding sequence. We obtained four additional mutations using ethyl methanesulfonate mutagenesis: three splicing junction mutations (*u851*, *u852*, and *u853*) and one missense mutation (*u854*; Figures S1A and S2A). Two *poml-1* alleles (*ok2266* and *tm4234;* Figures S1A and S2A) with the gene partially deleted were previously known. For many of the experiments we used the *ok2266* allele, which acted as a null, since it produced the same phenotype as *poml-1(u881;* see below).

None of the eight mutations produced touch insensitivity or any other obvious phenotype. The mutations, however, did render animals containing sensitizing mutations touch insensitive (Figures 2A and S2B). These sensitizing mutations were temperature-sensitive alleles of *mec-4* and *mec-6* (Gu *et al*., 1996), two hypomorphic alleles of *mec-6* [*u511*(G235E) and *u518*(G213E)] (García-Añoveros, 1995), and null alleles of *crt-1*, which encodes the ER chaperone calreticulin (Park *et al*., 2001; Xu *et al*., 2001). Except for the *crt-1* mutations, which lower MEC-4 protein level and cause slight touch defects (Xu *et al*., 2001), none of the sensitizing mutations produced touch insensitivity on their own. All the mutations produced severe touch insensitivity in *poml-1* null mutants. These synthetic phenotypes suggest a role for *poml-1* in TRN touch sensitivity. The loss of MEC-6, POML-1, or CRT-1 did not affect the general physiology of the TRNs, since light-activation of TRN-expressed channelrhodopsin-2 (Nagel *et al*., 2003) produced the same response in *crt-1*, *mec-6; poml-1*, and *crt-1; poml-1* mutants as in wild type (Figure S2C).

**Figure 2.**
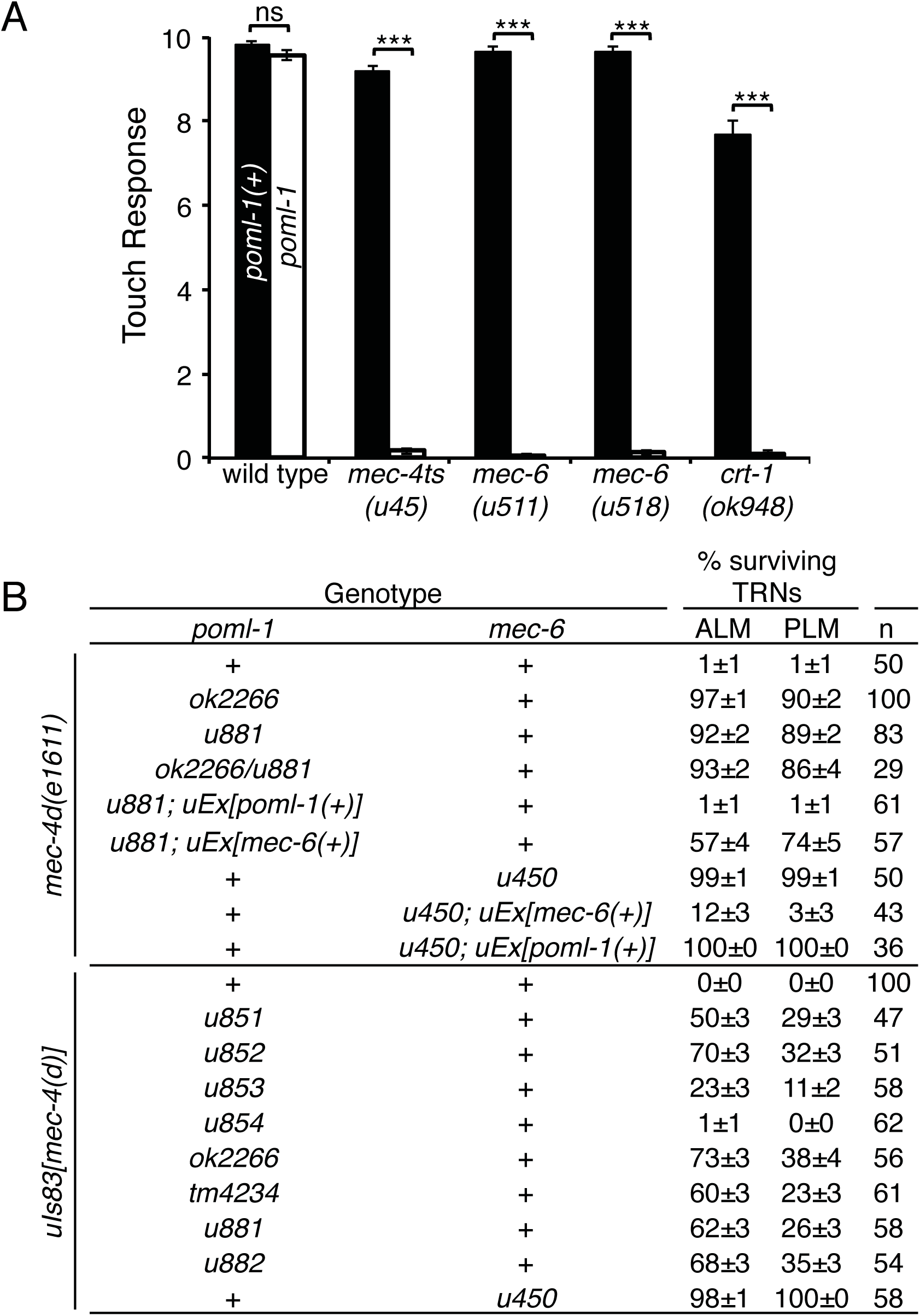
POML-1 is required for touch sensitivity and *mec-4(d)-*induced TRN degeneration. (A) *poml-1(u882)* reduced the touch response to 10 touches in sensitized backgrounds (30 animals were examined in three independent experiments). (B) *poml-1* and *mec-6* mutations suppress *mec-4*(*d*) degenerations. n indicates the number of animals examined. TRNs labeled with GFP in L4 and young adult animals were scored as having survived. *uIs83* is an integrated array that overexpresses *mec-4*(*d*). The rescue experiments used 3-5 stable lines.

Other evidence for a role of POML-1 in MEC-4 function comes from the suppression by *poml-1* mutations of the TRN degeneration caused by the hyperactive channel encoded by the *mec-4(d)* gain-of-function mutation *e1611* (Driscoll and Chalfie, 1991; Brown *et al*., 2007). *mec-6* mutations but not those of other touch sensitivity genes, suppressed these deaths (Chalfie and Wolinsky, 1990; Huang and Chalfie, 1994). In contrast to their weak effects on touch sensitivity, seven of the eight *poml-1* alleles (all but the missense mutation *u854*) strongly suppressed *mec-4(d)* neuronal degeneration to different extents (Figure 2B) and did so on their own. The suppressed animals were all touch insensitive, suggesting that either insufficient MEC-4(d) is available for touch sensitivity or that the mutant channels cannot transduce touch.

Overexpressing *poml-1(+)* rescued the *poml-1* phenotype in *poml-1 mec-4*(*d*) animals (resulting in TRN degeneration), but not the *mec-6* phenotype in *mec-6 mec-4*(*d*) animals (Figure 2B). Similarly, overexpressing *mec-6(+)* caused almost all the TRNs to die in *mec-6 mec-4*(*d*) animals, but only 35% of the TRNs in *poml-1 mec-4*(*d*) animals (Figure 2B). These results suggest that *mec-6* and *poml-1* have activities that cannot be replaced by the other gene.

Overexpressing *mec-4(d)* by the multiple copy insertion *uIs83* partially caused degeneration in *poml-1* animals, but not in animals with the *mec-6(u450)* mutation (Figure 2B), which deletes most of *mec-6* coding sequence and is considered to be a null allele (Chelur *et al*., 2002). Unlike *mec-6* mutations (Chalfie and Wolinsky, 1990; Shreffler *et al*., 1995), *poml-1* mutations did not suppress other DEG/ENaC gain-of-function mutations, including *deg-1* (Figure S2D) and *unc-8* (20 out of 20 animals were still Unc), even though POML-1 and DEG-1 are both expressed in IL1 neurons.

### POML-1 modulates MEC-4(d) channel activity in *Xenopus* oocytes

As with MEC-6 (Chelur *et al*., 2002), POML-1 increased the activity of the MEC-4(d) channel in *Xenopus* oocytes (Figures 3A and S3A). POML-1 increased the amiloride-sensitive Na^+^ current by three-fold, a smaller effect compared to the 10-fold increase produced by MEC-6. In addition, N-terminally EGFP-tagged POML-1 and N-terminally Myc-tagged MEC-4(d) immunoprecipitated each other in oocytes (Figure 3B). Moreover, POML-1 also immunoprecipitated C-terminally HA-tagged MEC-6 (Figure 3B), which is consistent with their colocalization in the ER and suggests they physically interact. Like MEC-6, POML-1 worked synergistically with MEC-2, but not with MEC-6, to increase channel activity ~40-fold (Figure 3A). [MEC-2, a stomatin-like protein that binds cholesterol, increases MEC-4 channel activity both in vivo and in vitro, perhaps through modulating the lipid environment surrounding the channel (Goodman *et al*., 2002; O’Hagan *et al*., 2005; Huber *et al*., 2006)]. At higher concentrations POML-1 on its own, like MEC-6 (Chelur *et al*., 2002), produced an amiloride-resistant current in oocytes (Figure S3, B and C).

**Figure 3.**
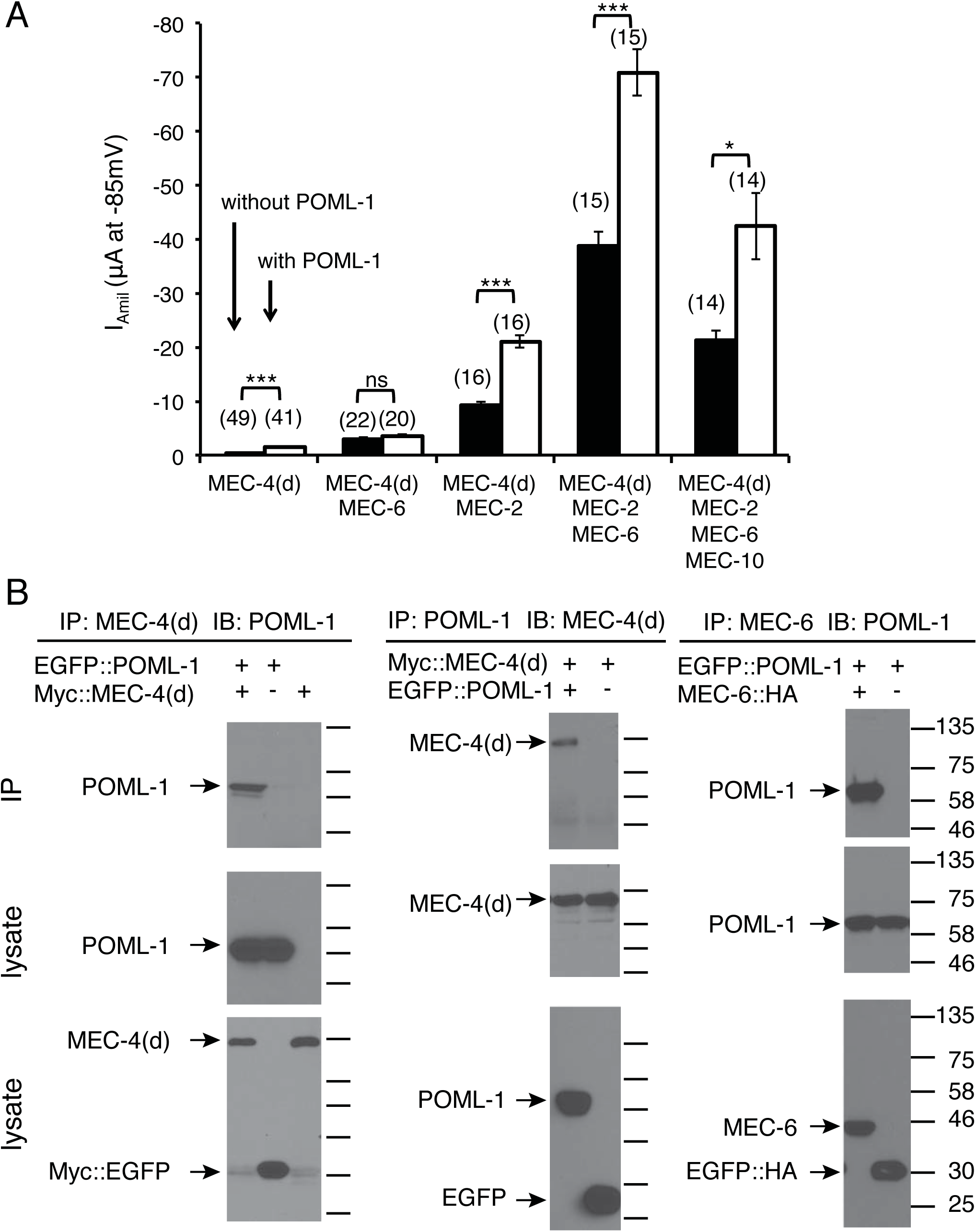
The effect of POML-1 on MEC-4(d) channel activity and their physical interaction in *Xenopus* oocytes. (A) Effect of POML-1 (white bars) on the MEC-4(d) amiloride-sensitive current at −85 mV in oocytes. The number of oocytes tested is given in parentheses. The oocytes were from at least two frogs. (B) Immunoprecipitation of POML-1 with MEC-4(d) and MEC-6 expressed in oocytes. Images are representative of 2-3 independent experiments. Molecular weights (kDa) of the protein markers are indicated on the right. EGFP::POML-1 is functional since its co-expression with MEC-4(d) generated amiloride-sensitive currents (EGFP::POML-1 and Myc::MEC-4(d), I_amil_ =-1.4±0.3 μA, n=5) that were similar to those of co-expressing untagged POML-1 and MEC-4(d) (I_amil_=-1.6±0.6 μA, n=5) 6 days after cRNA injection. The negative control (-) is EGFP with the same tag.

### POML-1 and MEC-6 affect the amount and distribution of MEC-4

Since POML-1 and MEC-6 affected MEC-4 channel function in vivo and channel activity in oocytes (Figures 2 and 3), we next tested their effect on the production and distribution of MEC-4. We examined wild type MEC-4 expression in cultured TRN cells with anti-MEC-4 antibodies. Both total expression (detected in permeable cells) and surface expression (detected in intact cells) of MEC-4 were substantially reduced by *mec-6* and *poml-1* mutations. MEC-4 surface expression was barely detected in cells with a *mec-6* null mutation and was reduced by 50% in cells with a *poml-1* null mutation (Figure 4, A and B). MEC-4 partially colocalized with ER and endosome markers in wild type TRN cell bodies but not with Golgi markers (Figure 4C). Less MEC-4 protein appeared in the TRN cell bodies in *mec-6* and *poml-1* mutants, but the localization of MEC-4 vis-à-vis the ER, endosome, and Golgi markers did not dramatically change (Figure 4, C and D).

**Figure 4.**
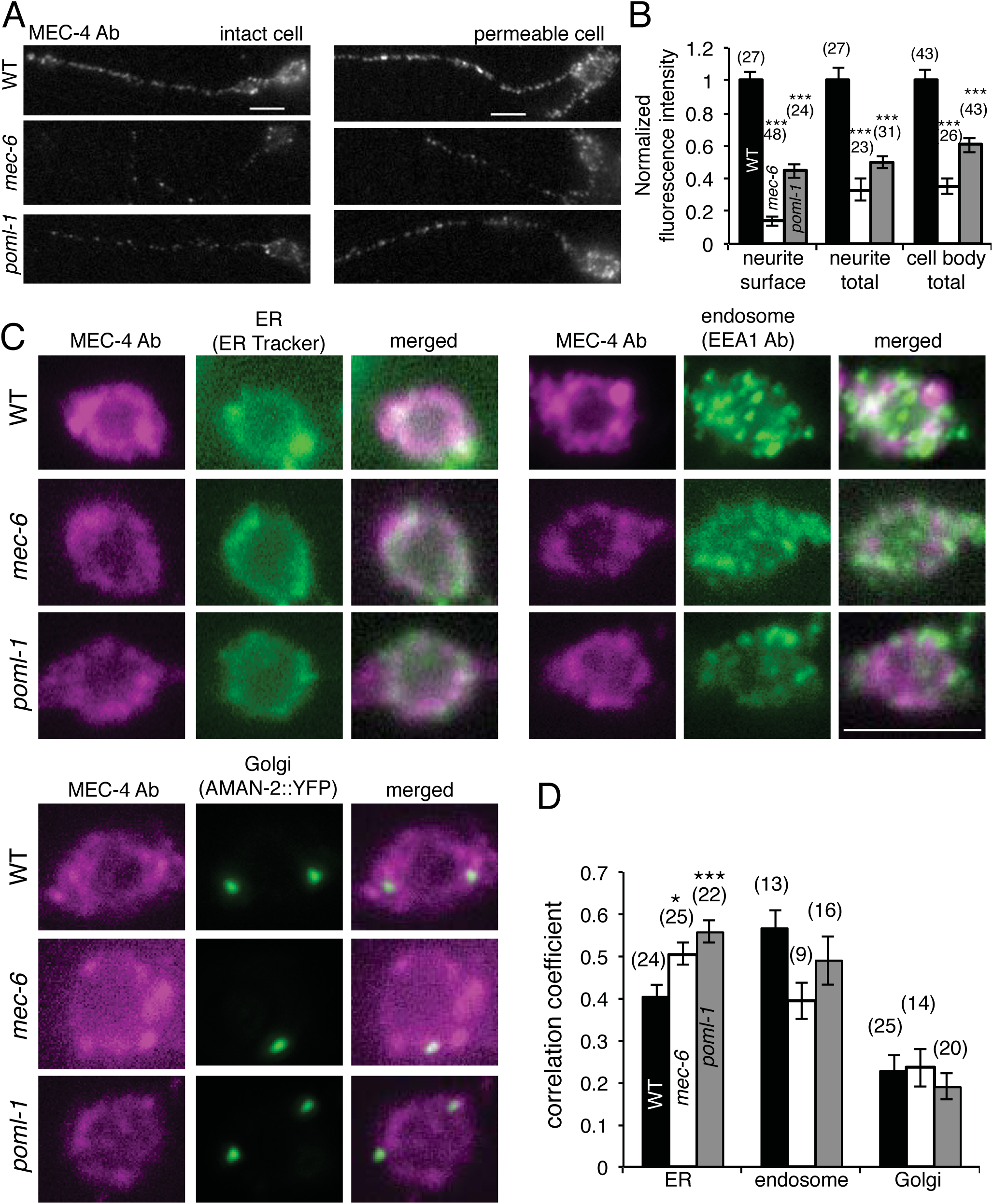
MEC-4 expression in cultured TRNs with *mec-6(u450)* or *poml-1(ok2266)* mutation. The images (A) and quantification (B) of MEC-4 expression as detected by an anti-MEC-4 antibody in intact (left panel) and permeabilized (right panel) TRNs in culture. MEC-4 immunofluorescence intensity was normalized to that of wild type (WT) TRNs. Scale bars = 5 μm (here and in C). Because most cell bodies leaked after immunostaining for intact cells (as evident by the staining for MEC-18, a cytoplasmic protein), immunofluorescence was only measured for MEC-4 surface expression in intact neurites (which did not show MEC-18 staining; Materials and Methods). Statistical significance is indicated for comparison with the wild type cells by one-way ANOVA with Tukey post hoc (here and D). The number of cell bodies tested from 2-3 experiments is given in parentheses (here and D). The subcellular localization of MEC-4 and markers for the ER, endosome, and Golgi (C) in cultured TRNs and their correlation coefficient (D).

We also examined MEC-4 expression in vivo using a fluorescent protein tag and an anti-MEC-4 antibody. Tagged MEC-4, which produced very strong fluorescence, appeared as large spots in the TRN cell body and smaller puncta in the neurite (Figure 5A; in some cases the spots were less prominent in the cell body and a meshwork was seen). Such MEC-4 aggregates were also detected by the anti-MEC-4 antibody in cell bodies, though this fluorescence was much less bright (under these conditions the diffuse expression was more obvious). In the cell body, MEC-4::TagRFP spots partially colocalized with POML-1 (Figure 5B) and the ER (YFP::TRAM-1), but some were always adjacent to the Golgi (AMAN-2::YFP, n= 20 TRNs) and the ER exit site (SEC-23::GFP; n = 10 TRNs, Figure S4A). Thus, MEC-4 may reside, at least for some time, in ER-Golgi intermediate compartment (Appenzeller-Herzog and Hauri, 2006) or trans-Golgi network (Traub and Kornfeld, 1997).

**Figure 5.**
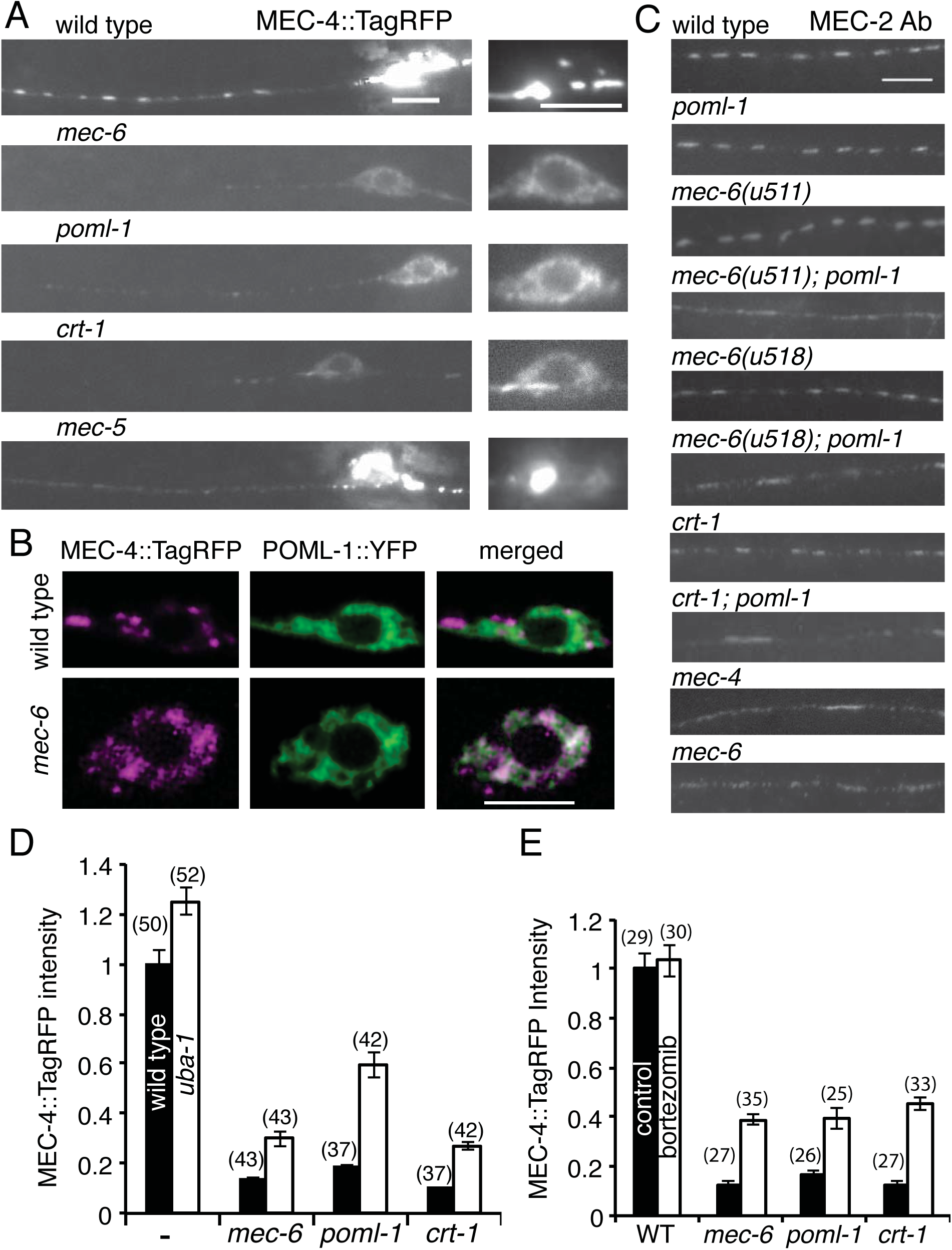
Effect of *mec-6*, *poml-1*, and *crt-1* mutations on MEC-4 and MEC-2 expression in the TRNs. Unless noted the following mutations were used: *crt-1(ok948)*, *mec-4(u253)*, *mec-5(u444)*, *mec-6(u450)*, *uba-1(it129)*, and *poml-1(ok2266)*. (A) MEC-4::TagRFP expression. Left panels show the merged images of expression at ten focal planes; right panels are images of the single plane showing the best focused image of the same cell body. Scale bar (here and in B and C) = 5 μm. (B) Confocal images of MEC-4::TagRFP and POML-1::YFP in the TRN cell body of wild type animals (upper panel) and *mec-6(u450)* mutants (lower panel). Their correlation coefficient is 0.2 ±0.04 and 0.7 ± 0.04 in wild type and *mec-6* mutants (n=10 TRNs), respectively. (C) MEC-2 expression in the TRN neurite. Images were representative of 20-30 TRNs examined in two experiments. (D) MEC-4::TagRFP fluorescence intensity (normalized to wild type) in PLM cell bodies of L4 larvae and young adults of controls, *mec-6(u450)*, *poml-1* and *crt-1* without (black) or with (white) a *uba-1* mutation. For each pair of white and black bars, the effect of the *uba-1* mutation was significant at p<0.001, except for wild type, which was p<0.01. The comparison to the control within each group (with or without *uba-1*) was significant at p<0.001 by two-way ANOVA with Bonferroni post-test. The number of examined PLM cells collected from three independent experiments is given in parentheses (here and E). (E) Fluorescence intensity of MEC-4::TagRFP (normalize to the wild type control) in PLM cell bodies of wild type (WT) and mutants either untreated (black) or treated (white) with 50 μM bortezomib for 8 hrs. Statistical comparisons are as in (D) with the exception that the difference between treated and untreated wild type animals was not significant.

Consistent with the results seen in cultured TRNs, *mec-6* and *poml-1* mutations reduced MEC-4 protein levels in vivo as seen with fusion proteins (Figure 5A) and an anti-MEC-4 antibody (Figure S4B). As in cultured TRNs (Figure 4, A-B), the reduction, was stronger with *mec-6* in vivo (Figure S4, B and C), and, thus, correlates with the stronger phenotype of *mec-6* with regard to touch sensitivity and *mec-4(d)* degeneration. Proteins were mainly found in the perinuclear mesh-like structure in the TRN cell body, indicating the ER, and puncta were not seen in the neurites. Indeed, in the TRN cell body with *mec-6* mutations, MEC-4 fusion proteins colocalized with POML-1 in the ER (Figure 5B). In addition, the half maximal pressure (P_1/2_) for the mechanoreceptor current (MRC) differed slightly between *poml-1* and wild type TRNs (*poml-1* P_1/2_ = 7.7 ± 0.8 nN/μm^2^ versus wild type P_1/2_ = 4.5 ± 0.7 nN/μm^2^, mean ± SD, Figure S4D), but *poml-1* mutations did not affect MRC peak amplitude or kinetics (Figure S4E). The effects of *mec-6* and *poml-1* mutations on MEC-4 appeared to be specific, because they did not affect the expression of each other (Figure S5A) or MEC-18 (Figure S4B).

The effect of *mec-6* and *poml-1* mutations was similar to that of a *crt-1* null mutation, which also reduced the amount of MEC-4 and its appearance as puncta in TRN cell bodies and neurites (Figure 5A). We found that MEC-6, POML-1, and CRT-1 affected the expression and distribution of MEC-4 protein, rather than the amount of *mec-4* mRNA as detected by single molecule fluorescence *in situ* hybridization (*crt-1* mutation slightly increased the number of *mec-4* mRNA molecules in TRNs, Figure S5B).

The effects of *mec-6* and *poml-1* mutations on MEC-4 protein levels and distribution suggest that MEC-6 and POML-1 act early in MEC-4 production and/or transport. Consistent with this hypothesis, mutations that presumably affect touch sensitivity after MEC-4 is made [e.g., null mutation of *mec-5*, a gene encoding an ECM collagen needed for touch sensitivity (Emtage *et al*., 2004)], disrupted the neurite localization of MEC-4 without affecting the level of MEC-4 protein (normalized intensity of MEC-4::TagRFP in the cell body, wild type 1±0.06 versus *mec-5(u444)* 0.92±0.05, n=29 PLM cells, not significant by the Student’s t test) or its distribution in the cell body (Figure 5A).

Since the production of MEC-2 puncta requires MEC-4 (Emtage *et al*., 2004; Zhang *et al*., 2004), we also tested the role of POML-1 on MEC-2 distribution using an anti-MEC-2 antibody in *poml-1* mutants, the two *mec-6* hypomorphic mutants *u511*(G235E) and *u518*(G213E), *crt-1* mutants, and *mec-6 (u511); poml-1, mec-6 (u518); poml-1* and *crt-1; poml-1* double mutants. No single mutation caused obvious defects in MEC-2 puncta. In contrast, the double mutants, as in the null mutants of *mec-4* and *mec-6*, had disrupted MEC-2 puncta (Figure 5C), a result that is consistent with the need for the double mutations to cause touch insensitivity.

### MEC-6 and POML-1 likely act as chaperones

The similarity of the phenotypes of *crt-1*, *mec-6*, and *poml-1* mutants, the expression of all three proteins primarily in the ER, and the additivity of their phenotypes with regard to touch sensitivity suggest that MEC-6 and POML-1, like CRT-1, may, at least in part, act as chaperones. If these proteins facilitate the folding/assembly of MEC-4 in the ER, the reduction in MEC-4 protein in these mutants, could be a consequence of increased protein degradation. Indeed, the loss of CRT-1, MEC-6, or POML-1 caused an increase in MEC-4 degradation. Mutation of the ubiquitin-activating (E1) enzyme gene *uba-1* (Jones *et al*., 2002) or treatment of animals with the proteasome inhibitor bortezomib increased MEC-4 levels 2-3 fold in *mec-6*, *poml-1*, and *crt-1* mutants (Figure 5, D and E). *uba-1* mutation, but not bortezomib, affected the removal of wild type MEC-4 from the cell surface (Chen and Chalfie, 2015). The effects of bortezomib on MEC-4 levels in the *mec-6*, *poml-1*, and *crt-1* mutants, thus, reveal a different process, perhaps a consequence of the accumulation of misfolded protein.

This increase, however, did not restore MEC-4 levels to those seen in wild type, in part, perhaps, because less MEC-4 protein was produced in *mec-6*, *poml-1*, and *crt-1* mutants even when the degradation pathway was blocked or the *uba-1* mutation and bortezomib treatment only partially suppressed the degradation pathway. Although these treatments increased the amount of MEC-4, they did not change its distribution. MEC-4 was still largely restricted to the same mesh-like structure at perinuclear sites in the cell body seen in the mutants without the *uba-1* mutation or bortezomib treatment (Figures 5A and S5C). Moreover, the MEC-4 puncta were not restored in TRN neurites (Figure S5C). Additionally, because *uba-1* mutation did not restore touch sensitivity to *mec-6* null mutants or *crt-1; poml-1* double mutants (Figure S5D) and only resulted in a modest increase (16% in ALM for *uba-1*) of touch cell deaths in *poml-1 mec-4(d)* animals but not in *mec-6; mec-4*(*d*) animals (Figure S5E), the increased MEC-4 does not function, perhaps because it is misfolded or not properly localized.

Since overexpression of the ER transport protein SEC-24 rescued the trafficking defects caused by the loss of a putative ER chaperone/cargo receptor but not the folding defects in yeast (Pagant *et al*., 2015), we tested whether overexpression of *C. elegans sec-24* genes (*sec-24.1* and *sec-24.2*) in the TRNs could similarly suppress the effects of *crt-1*, *mec-6*, and *poml-1* mutations. [CRT-1 is also required for the degeneration caused by hyperactive MEC-4(d) channels (Xu *et al*., 2001)]. The effects were partial: about 50% of the TRNs died in *poml-1 mec-4(d)* animals and 30% in *crt-1; mec-4(d)* animals, but no TRNs died in *mec-6; mec-4(d)* animals overexpressing the *sec-24* genes (Figure 6A). [We also noticed that overexpression of *sec-24(+)* caused morphological defects: wavy neurites, extra neurites and misplaced cell bodies; these defects were rarely seen in animals that did not overexpress *sec-24(+)*(Figure S5F).] In addition the overexpression of the *C. elegans sec-24* genes with mutated potential cargo-binding sites [corresponding to Yeast SEC-24 R230A, R235A, L616W (Miller *et al*., 2003)] reduced but did not eliminate the effect; 20% of the TRNs died in *poml-1 mec-4*(*d*) animals (Figure 6A). Although overexpressing SEC-24 in *poml-1* mutants doubled the amount of MEC-4::TagRFP in proximal TRN neurites, but not in the cell bodies (Figure 6B), it did not restore touch sensitivity to *mec-6(u511); poml-1* animals (0.21±0.09 response to 10 touches, n=20 animals).

**Figure 6.**
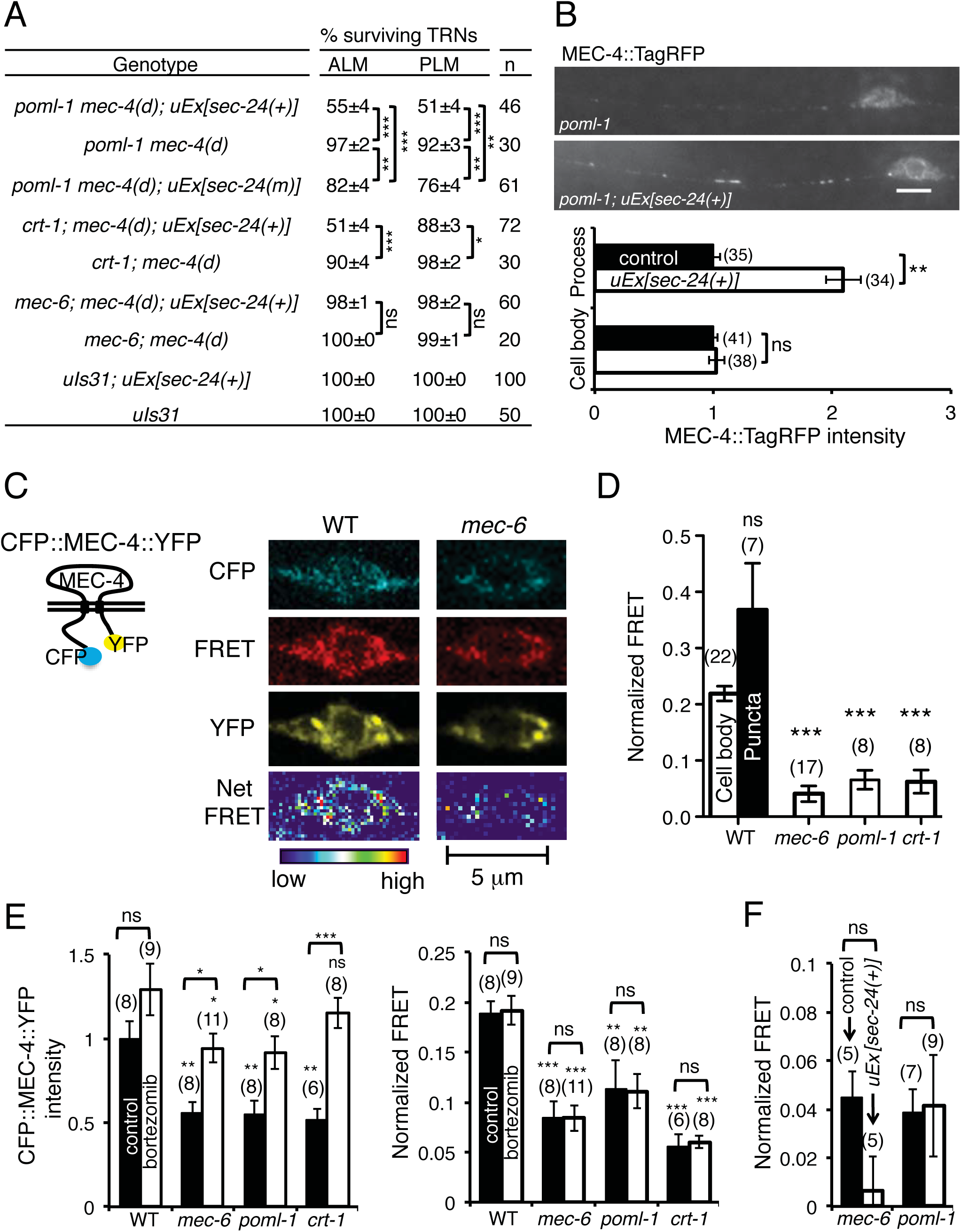
MEC-6, POML-1 and CRT-1 may function as chaperones. The following mutations were used: *mec-6(u450)*, *poml-1(ok2266)* and *crt-1(ok948)*. (A) The effect of *sec-24.1* and *sec-24.2* overexpression, *uEx*[*sec-24(+)*], on the suppression of *mec-4*(*d*) deaths by *poml-1*, *crt-1*, and *mec-6* mutations. In some strains the cargo-binding sites of *sec-24.1* and *sec-24.2* were mutated [*sec-24(m)*]. n = the number of animals examined. The results with *uEx[sec-24(+)] and uEx[sec-24(m)]* were collected from 2-5 stable lines. (B) Effect of overexpressing *sec-24(+)* on MEC-4::TagRFP fluorescence intensity in the PLM cell bodies and proximal neurites of L4 larvae and young adults with the *poml-1* mutation. Data with *uEx[sec-24(+)]* were collected from two stable lines. The number of PLM cells examined is given in parentheses. Fluorescence intensity was normalized to that of PLM in *poml-1* mutants (control). Scale bar = 5 μm. (C) Schematic of CFP::MEC-4::YFP protein (left panel) and images of CFP::MEC-4::YFP in the TRN cell body taken with the CFP (blue), FRET (red), and YFP (yellow) channels, respectively (right panel). The Net FRET signal is given by a pseudocolored image to show the relative intensity. (D) The normalized FRET signal (see Materials and Methods) of CFP::MEC-4::YFP either in the TRN cell bodies (white bars) of wild type animals (WT) and mutants or in the puncta of wild type animals (black bar). The number of cell bodies or strongly fluorescent puncta tested (from 2-3 stable lines with extrachromosomal arrays collected from 3-4 experiments) is given in parentheses. Statistical significance is indicated for comparison with the FRET signal in the wild type cell body. (E) CFP::MEC-4::YFP intensity (measured in the YFP channel and normalized to that of wild type controls) and FRET signals in the TRN cell body of wild type, *mec-6*, *poml-1*, and *crt-1* animals treated with bortezomib. The number of examined cells bodies here and in F is indicated in parenthesis. These experiments used cells from an integrated line, which produced similar FRET signals to those stable lines with extrachromosomal arrays that were used in panel D. Bortezomib treatment had a significant effect on CFP::MEC-4::YFP intensity [left panel, F(1, 58)=33.56, p<0.0001], but no effect on the FRET signal [right panel, F(1, 58)=0.01, p=0.9162, by two-way ANOVA with Bonferroni post-tests]. The value above the bracket is that of the pairwise comparison. The values below the bracket are for the comparison to the wild type (WT) of each untreated (black bars) or treated group (white bars) by two-way ANOVA with Bonferroni post-tests. The difference in CFP::MEC-4::YFP fluorescence intensity (left panel) between control wild type and bortezomib-treated *mec-6*, *poml-1*, and *crt-1* mutants was not significant. (F) FRET signals in *mec-6* and *poml-1* animals overexpressing *sec-24(+)* in TRNs.

These results can be explained if little, if any, MEC-4 folds in *mec-6* mutants, but some MEC-4 folds but is not transported to the surface in *poml-1* and *crt-1* mutants. Thus, by increasing transport to the surface, SEC-24 could cause more *mec-4(d)*-induced deaths in the *poml-1* and *crt-1* mutants but not in the *mec-6* mutants because they contain no functioning protein. These results suggest an absolute need for MEC-6 in MEC-4 expression and distribution and are consistent with the need for MEC-6 (Chalfie and Sulston, 1981), but not of POML-1 and CRT-1, in touch sensitivity in wild type animals (Figure 2A).

CRT-1 can bind Ca^2+^ and regulate Ca^2+^ homeostasis in the ER (Michalak *et al*., 2009). Xu *et al*. (2001) have suggested that *crt-1* suppression of MEC-4(d)-induced cell death is attributed to the Ca^2+^ binding capacity of CRT-1 in the ER and can be partially reversed by the release of ER Ca^2+^ induced by thapsigargin. We tested whether *poml-1* and *mec-6* suppress MEC-4(d) through a similar mechanism, and found that, in contrast to *crt-1*, the effect of *poml-1* and *mec-6* on cell death was not affected by thapsigargin treatment (Figure S5G). Thus, *poml-1* and *mec-6* mutations are less likely to suppress MEC-4(d)-induced cell death through affecting the subcellular Ca^2+^ level. Moreover, manipulating subcellular Ca^2+^ level has no effect on MEC-4 expression (Xu *et al*., 2001). Therefore, *mec-6* and *poml-1* suppression of MEC-4(d) is primarily due to their effect on MEC-4 expression and distribution.

Because Förster resonance energy transfer (FRET) can be used to monitor protein folding (Philipps *et al*., 2003), we used a CFP::MEC-4::YFP fusion to examine whether MEC-6, POML-1, and CRT-1 affect the MEC-4 protein folding. This fusion was expressed in the TRNs: as with MEC-4::GFP and MEC-4::TagRFP, the protein formed regular puncta in the neurite (Figure S5H) and a mesh-like structure and spots in the cell body, though the spots in the cell body were smaller and dimmer than with MEC-4::TagRFP (Figure 6C). In *mec-6, poml-1*, or *crt-1* mutants, CFP::MEC-4::YFP was restricted to the cell body, where the fluorescence intensity was reduced by nearly 50% (Figure 6C). CFP::MEC-4::YFP produced an efficient FRET signal in wild type animals (Figure 6, C and D), suggesting that CFP and YFP were close to each other.

In contrast to the FRET signal in wild type, FRET from CFP::MEC-4::YFP was reduced by 70-80% in *mec-6, poml-1*, or *crt-1* mutants (Figure 6, C and D). The reduction of FRET in these mutants was not due to reduced CFP::MEC-4::YFP expression, because the FRET signal was normalized to the CFP and YFP intensities (see Material and Methods) and wild type animals expressing similar level of CFP::MEC-4::YFP to these mutants [from *uIs190(mec-4p::cfp::mec-4::yfp)/+* animals] produced robust FRET signals (normalized FRET: 0.24 ± 0.06, n=5). Moreover, bortezomib treatment increased CFP::MEC-4::YFP expression in *mec-6, poml-1*, and *crt-1* mutants to a level similar to that in wild type untreated animals, but had no effect on the FRET signal (Figure 6E). These data suggest that the folding and/or assembly of MEC-4 is compromised in these mutants. Overexpression of SEC-24 also failed to increase the FRET signals in *mec-6* and *poml-1* mutants (Figure 6F), a result that is consistent with the role for SEC-24 in protein transport but not in protein folding.

Given the localization of MEC-6 and POML-1 in the ER and their potential effects on folding, we wondered whether *mec-6* or *poml-1* mutations induced a general ER stress response. These mutations, however, did not significantly change the expression of *xbp-1b::gfp* (Figure S5I), which produces a GFP translation product only in response to ER stress (Shim et al., 2004). Wild type and the mutant strains showed a similar ER stress response, when proteasomes were inhibited by bortezomib (Figure S5I).

### MEC-6 accelerates the surface expression of MEC-4 in *Xenopus* oocytes

Consistent with its role as a chaperone, we found that MEC-6 greatly increased the amount of surface expression of MEC-4 in *Xenopus* oocytes using total internal reflection (TIRF) microscopy and biotinylation (Figure 7, A-D) two days after cRNA injection. In contrast, MEC-2 and POML-1 did not increase the surface expression of MEC-4 (Figure 7, A-D). Although TIRF microscopy cannot be used to determine the position of the protein on the plasma membrane, the biotinylation experiments suggest that MEC-6 affects MEC-4 surface expression. These results support the hypothesis that MEC-6 assists the maturation of MEC-4 channels, and/or its transport to the plasma membrane.

**Figure 7.**
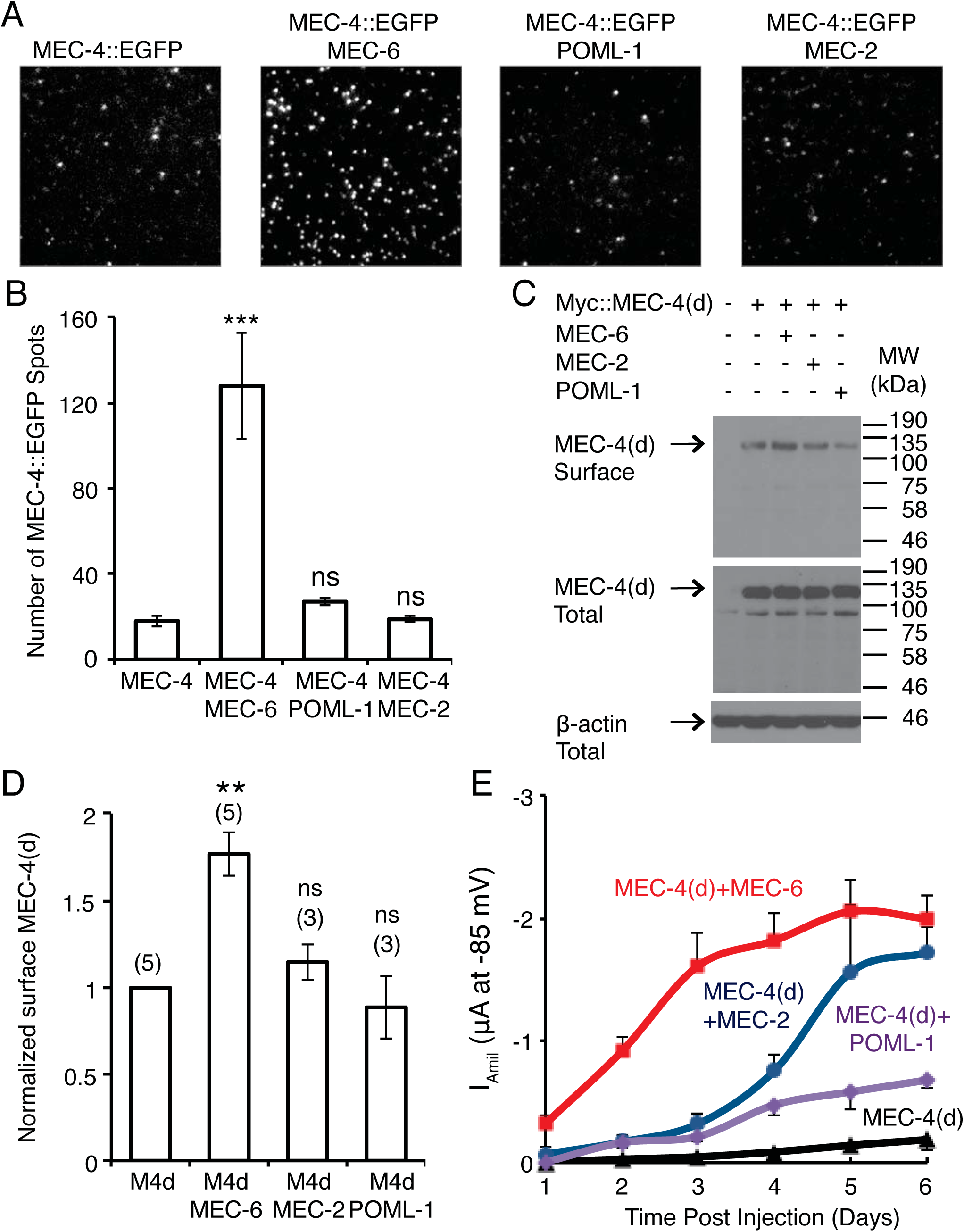
The effect of MEC-6 and POML-1 on MEC-4 surface expression in *Xenopus* oocytes. Images (A) and quantification (B) of MEC-4::EGFP fluorescent spots on the oocytes surface visualized by TIRF imaging (n = 19-29 patches from 14-16 cells from two different batches) 2 days after cRNA injection. Values are compared to the expression of MEC-4::EGFP alone using the Mann-Whitney test. The field dimensions are 13 μm x 13 μm. (C) Western blot of Myc::MEC-4(d) on the surface of oocyte as detected by biotinylation (top) and the expression of Myc::MEC-4(d) in total lysate of oocytes (middle) at 2 days after cRNA injection. β-actin detected in total lysate was used as an input control (bottom). Molecular weights (kDa) of the protein markers are indicated on the right. (D) Quantification of changes in surface Myc::MEC-4(d) detected by biotinylation at 2 days after cRNA injection (the number of independent experiments is given in parenthesis). All data are normalized and compared to Myc::MEC-4(d) expression alone by the one sample t test. MEC-6, MEC-2 and POML-1 did not affect Myc::MEC-4(d) levels in total lysates at 2 days after injection (MEC-6 1.0±0.1, MEC-2 1.0±0.1, POML-1 0.9±0.1, n=4-5 independent experiments, normalized and compared to the expression of Myc::MEC-4(d) alone, not significant by one sample t test). The normalized amount of total Myc::MEC-4(d) differed by no more than 25% in any of the experiments. (E) The amiloride-sensitive MEC-4(d) current at −85 mV [12-18 oocytes (2 days after cRNA injection) or 6-12 oocytes (other times) of three batches] on its own and in the presence of MEC-2, MEC-6, and POML-1 at various times after cRNA injection. p<0.001 for I _amil_ at −85 mV between oocytes 2 days after injected with MEC-4(d) and MEC-6 vs MEC-4(d) alone, MEC-4(d) and MEC-2, or MEC-4(d) and POML-1; no statistically significance was found between oocytes 2 days after injected with MEC-4(d) alone vs MEC-4(d) and MEC-2, or MEC-4(d) and POML-1. p<0.001 for I _amil_ at −85 mV between oocytes 1 day after injected with MEC-4(d) and MEC-6 vs MEC-4(d) alone; p<0.01 between MEC-4(d) and MEC-6 vs MEC-4(d) and MEC-2, or MEC-4(d) and POML-1. One-way ANOVA with Tukey post hoc.

The effect of MEC-6 on MEC-4(d) surface expression in oocytes was not seen five days after injection by biotinylation (Figure S6A), which is consistent with the previous experiments (Chelur *et al*., 2002; Brown *et al*., 2008). Presumably, the maximum steady state amount of MEC-4 is found on the surface with or without MEC-6 by five days. The lack of an effect in those previous experiments is due, at least in part, to the longer period of expression, and presumably to the accumulation of more inactive MEC-4 in the absence of MEC-6.

We also tested whether the early, MEC-6-induced change in membrane-associated MEC-4 affected the MEC-4(d) current in oocytes. Indeed, MEC-6, but not MEC-2 or POML-1, increased the MEC-4(d) current over 30-fold to nearly 50% of the maximum current two days after injection [Figure 7E; the fold difference is greater here than above (Figure 3A) because these oocytes had been injected with the lesser amount of *mec-4* cRNA, so the MEC-4(d) current was lower]. The early effect of MEC-6 compared to POML-1 and MEC-2 on MEC-4(d) activity two days after injection was due to different amounts of surface MEC-4(d) rather than of total MEC-4(d) (Figure 7, C and D). Differences in the timing of MEC-6, MEC-2 and POML-1 expression were unlikely to cause these differences in the MEC-4(d) current. MEC-6, MEC-2 and POML-1 were all expressed well two days after injection, and all had higher expression level five days after injection (Figure S6, B-D). In addition, although injecting oocytes with greater amounts of *mec-2* and *poml-1* cRNA increased the levels of total MEC-2 and POML-1 at two days after injection to the levels normally seen at five days after injection (Figure S6, C and D), MEC-4(d) currents two days after injection were not increased (Figure S6, E and F).

In addition to MEC-6 increasing MEC-4 membrane expression, it also increased, albeit weakly, the membrane expression of MEC-2 in oocytes (Figure S6, G and H). In contrast, although co-expression of POML-1 with MEC-2 doubled MEC-4(d) channel activity over that in oocytes without POML-1 (Figure 3A), POML-1 did not affect MEC-2 surface expression (Figure S6, G and H). In addition neither POML-1 nor MEC-6 changed total MEC-2 expression in oocytes (Figure S6I), stabilized MEC-2 (which moves on the surface of oocytes; Chen *et al*., 2015), or caused it to colocalize with MEC-4 (Video S1).

## DISCUSSION

MEC-6 is essential for TRN touch sensitivity (Chalfie and Sulston, 1981; Chelur *et al*., 2002). Previously the interaction of MEC-4 and MEC-6 (Chelur *et al*., 2002) led us to conclude that MEC-6 was a component of the transduction channel. The finding that MEC-4 and MEC-6 do not colocalize on the plasma membrane of *Xenopus* oocytes (Chen *et al*., 2015) and that MEC-6 and the related protein POML-1 fail to localize with MEC-4 and MEC-2 in TRN neurites (this paper) argues against MEC-6 and POML-1 being components of the transduction complex. The primary localization of these proteins in the ER, their colocalization with MEC-4 in the ER, and their effect on the expression and localization of MEC-4, suggests that MEC-6 and POML-1 act instead as ER-resident chaperones. Thus, MEC-6 and POML-1 represent a new class of chaperones. The effects on functional expression of MEC-6 and POML-1 (and in the case of MEC-6 on transport) may underlie the increases in MEC-4(d) channel currents we saw in oocytes. MEC-6 and POML-1, however, are likely to affect relatively few proteins since general ER stress was not induced in *mec-6* or *poml-1* mutants

Although both MEC-6 and POML-1 are required for MEC-4 expression and localization, they do not act identically. Specifically, MEC-6, but not POML-1, accelerated the appearance of MEC-4 on the oocyte surface, and SEC-24 proteins partially suppressed the inhibition of *mec-4(d)*-induced TRN cell death caused by *poml-1*, but not *mec-6* mutations. These results and the finding that overexpression of *mec-6* or *poml-1* did not rescue mutations in the other gene argue that MEC-6 and POML-1 may have distinct but overlapping roles in MEC-4 channel maturation. In contrast to *mec-6*, *poml-1* mutants still have nearly normal touch sensitivity and MEC-2 puncta, indicating these animals have a reduced, but still sufficient, amount of functional MEC-4 in the TRN neurites. This MEC-4 can be detected on the surface of the cultured TRNs but does not form visible puncta in vivo, although they presumably are sufficient enough to allow for the formation of MEC-2 puncta, which depend on MEC-4 (Zhang *et al*., 2004). This result agrees with stoichiometry data suggesting that visible MEC-4 puncta are not necessary for the channel function (Chen *et al*., 2015).

In contrast to CRT-1, which has more general functions in cells, e.g., facilitating glycoprotein folding and regulating Ca^2+^ homeostasis in the ER (Michalak *et al*., 2009), the action of MEC-6 and POML-1 is more restricted. Unlike calreticulin, which can bind up to 25 Ca^2+^ ions (Baksh and Michalak, 1991; Michalak *et al*., 2009), MEC-6 and POML-1 lack a similar Ca^2+^ binding domain or even the two Ca^2+^ binding residues in mammalian PONs (Harel *et al*., 2004), and, thus, are unlikely to play a direct role in regulating subcellular Ca^2+^ content. Consistent with the absence of Ca^2+^ binding domains in MEC-6 and POML-1, induction of ER Ca^2+^ release by thapsigargin had no effect on the suppression of MEC-4(d) by *mec-6* and *poml-1* (such treatment does affect the suppression by *crt-1*; Xu *et al*., 2001). Moreover, genetic screens for mutations that reverse *poml-1* suppression of MEC-4(d) identified genes that encodes MEC-10 and MEC-19, which normally inhibit MEC-4(d) channel activity and surface expression (Chen *et al*., 2016). Therefore, the effect of *poml-1*, and perhaps *mec-6*, on MEC-4(d) is largely due to effects on MEC-4 expression and production, rather than modulation of cellular Ca^2+^.

Both MEC-6 and POML-1 are needed for the maturation of DEG/ENaC proteins. MEC-6, which is expressed in many neurons and muscles, is needed for the action of gain-of-function (d) mutations affecting several DEG-ENaC proteins (DEG-1 and UNC-8) and ectopically-expressed MEC-4(d), but not a gain-of-function, degeneration-causing mutation affecting the nicotinic acetylcholine receptor protein DEG-3 (Chalfie and Wolinsky, 1990; García-Añoveros, 1995; Shreffler *et al*., 1995; Harbinder *et al*., 1997).

The action of MEC-6 and POML-1 may not, however, be restricted to only DEG/ENaC proteins, proteins that form amiloride-sensitive channels. That both MEC-6 (Chelur *et al*., 2002) and POML-1 (this work) can produce an amiloride-resistant current in oocytes hints that they may act on other proteins. In addition, we found MEC-6, but not POML-1, increased MEC-2 surface expression by approximately 60% in oocytes. POML-1 might affect MEC-2 activity since POML-1 increased MEC-4(d) channel activity in oocytes with MEC-2 or with MEC-2 and MEC-6 together, but not with MEC-6 on its own. Since POML-1 did not change MEC-2 expression in oocytes or in the TRNs, we do not know whether this enhancement was direct, e.g., by altering MEC-2 conformation, or indirect through changes in MEC-4.

MEC-6 and POML-1 and the human PONs are ~27% identical (over the C-terminal 260 residues), and all have an N-terminal hydrophobic region (Sorenson *et al*., 1999). Interestingly, two of the three mammalian PONs, PON2 and PON3, are found in the ER and can reduce the cell death induced by the unfolded protein response (Horke *et al*., 2007; Schweikert *et al*., 2012). The characterization of MEC-6 and POML-1 in *C. elegans* suggests a novel function of this protein family: ER chaperones that facilitate the maturation and transport of DEG/ENaC and, perhaps in vertebrates, other proteins.

## MATERIALS AND METHODS

### *C. elegans* procedures

Unless otherwise indicated, strains were maintained and studied at 20°C on the OP50 strain of *E. coli* according to Brenner, 1974. All the translational fusions were based on pPD95.75 (www.addgene.org/static/cms/files/Vec95.pdf). Transgenic animals were prepared by microinjection and integrated transgenes were generated by UV irradiation (Chelur and Chalfie, 2007). Details about strains, plasmids, and cDNAs are given in the Supplemental Materials.

Ethyl methanesulphonate mutagenesis was performed according to described in Brenner, 1974 to obtain additional alleles of *poml-1* (See Supplemental Materials).

We studied gentle touch sensitivity in blind tests as described (Chalfie and Sulston, 1981). We quantified the response by counting the number of response to ten touches delivered alternately near the head and tail in 30 animals. We performed in vivo electrophysiology as previously described (O’Hagan *et al*., 2005). We also used blue light and channelrhodopsis-2 expressed in the TRNs to stimulate these cells as previously described (Chen and Chalfie, 2014).

We performed single molecule fluorescence *in situ* hybridization to count *mec-4* mRNA as previously described (Topalidou *et al*., 2011).

Bortezomib (Selleckchem, Houston, TX) was dissolved in DMSO to make 130 mM stock and added to NGM medium to make plates containing 50 μM bortezomib. L3-L4 stage larvae were transformed to NGM plates with bortezomib and grow for 8 hours. Animals get sick if treated for longer times. In the control group animals of the same age were transferred to NGM plates containing the same volume of DMSO without bortezomib.

### *C. elegans* microscopy and immunofluorescence

Confocal images were acquired using a 63X NA 1.40 oil immersion objective on a ZEISS LSM700 confocal microscope. Colocalization of MEC-6 and POML-1 with each other and with ER and Golgi markers in the cell body was quantified by the colocalization function in the ZEN 2010 software and is represented by Pearson’s correlation R value. Live animals were anesthetized using 100 mM 2-3 butanedione monoxime in 10 mM HEPES, pH 7.4.

Fluorescence was observed using a Zeiss Axio Observer Z1 inverted microscope equipped with a Photometrics CoolSnap HQ^2^ camera (Photometrics, Tucson, AZ) and a Zeiss Axioskop II equipped with a SPOT 2 slider camera (SPOT Imaging Solutions, Sterling Heights, MI).

Immunostaining of larvae and adults was performed according to Miller and Shakes, 1995, except for MEC-4 immunostaining, which used a mouse anti-MEC-4 antibody (ab22184, Abcam, Cambridge, MA) and was performed according to Bellanger *et al*., 2012. The following antibodies were used for immunostaining of *C. elegans*: anti-MEC-18 (Zhang, 2004), anti-MEC-2 N-terminus (Zhang, 2004), anti-MEC-4 (mouse, ab22184, Abcam), anti-FLAG (mouse, F1804, Sigma, St. Louis, MO), and anti-GFP (rabbit polyclonal A11122 and mouse monoclonal 3E6; Life Technologies, Carlsbad, CA) diluted 1:200, Rhodamine Red-X-conjugated goat anti-rabbit IgG, Alexa Fluor 488-conjugated goat anti-rabbit IgG, and Alexa Fluor 647-conjugated goat anti-rabbit/mouse IgG (Jackson ImmunoResearch Laboratories, West Grove, PA), and Alexa Fluor 488/555-conjugated goat anti-mouse (Life Technologies, Carlsbad, CA) diluted 1:700.

MEC-2 and POML-1 immunofluorescence puncta were analyzed over about 50 μm length of TRN neurites with regular puncta using Image J (rsbweb.nih.gov/ij/) and the Puncta Analysis Toolkit beta developed by Dr. Mei Zhen, Samuel Lunenfeld Research Institute, Toronto, Canada. The width of puncta was the length at half-maximum of each peak and the average distance between puncta were calculated as 1/(number of puncta per μm). Colocalization of POML-1, MEC-4, MEC-2 and MEC-6 puncta in the TRN neurite was analyzed by Image J plugin Coloc 2 (http://fiji.sc/Coloc2) as described in Chen *et al*., 2015.

We measured fluorescence intensity in the cell body by selecting the cell body area (20-30 μm^2^), and measuring the mean intensity subtracted from the background of the same size area by Image J. The intensity of the MEC-4::TagRFP puncta intensity was measured in the best focused image of six images taken at different z planes using the Puncta Analysis Toolkit beta. Puncta were examined over a region approximately equivalent to ten cell body lengths starting near the cell body.

Isolated, embryonic TRNs that had been cultured for 15-24 hrs (Zhang *et al*., 2002) were fixed in 4% paraformaldehyde, blocked in PBS with 1% BSA (in some experiments 0.2% Triton-X-100 was added to permeabilize the plasma membrane), incubated with primary antibodies (as indicated above) at 4°C for 2 hrs, washed 3X in PBS, incubated with secondary antibodies (Rhodamine Red-X-conjugated goat anti-rabbit IgG and Alexa Fluor 647-conjugated goat anti-mouse IgG diluted 1:2000) at room temperature for 30 mins, and washed 3X in PBS. Immunofluorescence of MEC-18, a cytoplasmic TRN protein, was used as an internal control for nonpermeabilized immunostaining (Chen and Chalfie, 2015). Most cell bodies and some neurites became leaky during immunostaining and displayed clear immunofluorescence signal of MEC-18. Only the fluorescence in intact neurites was measured. We quantified the mean immunofluorescence intensity of MEC-4 in the cell body and the neurite by Image J.

After immunostaining for MEC-4, cultured TRNs were incubated with ER-tracker Blue-White DPX (Life Technologies, E12353) diluted 1:1000 in PBS at room temperature for 30 mins and washed 3X in PBS prior to imaging. An anti-EEA1 antibody (ab2900, Abcam) diluted 1:400 were used to label endosome in cultured TRNs. Correlation coefficient of MEC-4 with markers for ER, endosome, and Golgi were analyzed using Image J plugin Coloc 2 (http://fiji.sc/Coloc2) as described above.

### FRET

FRET was performed on L4 to young adult animals glued with Dermabond (Ethicon Inc., Somerville, NJ) onto 2% agarose pads according to Youvan *et al*., 1997 using a ZEISS LSM700 confocal microscope. Mean fluorescence intensity minus background was determined in the cell body (and in the puncta for wild type) for three channels: CFP (CFP_excitation=405nm_ - CFP_emission=420-475nm_), YFP (YFPexcitation=488nm - YFP_emission≥520nm_), and FRET (CFPexcitation=405nm-YFP_emission≥520nm_). The cross talk of CFP into the FRET channel (Df = FRET/CFP = 84%) was determined in animals expressing *mec-4p::cfp::mec-4*. Similarly, the cross talk co-efficiency of YFP to the FRET channel is determined by expressing *mec-4p::mec-4::yfp* only, and calculated as Af = FRET/YFP = 1.4%. Net FRET was calculated as FRET – 0.84 X CFP – 0.014 X YFP (Youvan *et al*., 1997). Normalized FRET was calculated as Net FRET/(CFP X YFP)^1/2^ (Xia and Liu, 2001).

### Electrophysiology, biochemistry, and single molecule imaging in *Xenopus* ooctyes

A 1050 bp *poml-1* cDNA coding sequence was generated by RACE PCR using FirstChoice RLM-RACE kit (Ambion, Grand Island, NY) with mRNA from wild type animals, and cloned in pGEM-HE (Liman *et al*., 1992). cDNA of POML-1 was cloned into pGEMHE-EGFP-X (Ulbrich and Isacoff, 2007) to generate proteins tagged with EGFP at their N-termini.

cRNA expression and electrophysiology in *Xenopus laevis* oocytes (Xenopus I, Dexter, MI; Nasco, Fort Atkinson, WI; Ecocyte, Austin, TX) followed the procedures and used the plasmids previously described (Goodman *et al*., 2002). In the experiments described in Figures 3, 10 ng cRNA of MEC-4, MEC-2, MEC-10, 1 ng MEC-6 cRNA, and/or 5 ng cRNA of POML-1 were injected into oocytes unless noted. Oocytes were maintained as previously describe (Arnadóttir *et al*., 2011). Membrane current was measured 4-6 days after cRNA injection using a two-electrode voltage clamp as previously described (Goodman *et al*., 2002). In the experiments described in Figure7, A and B, oocytes were injected with 3.75 ng MEC-4::EGFP cRNA, 1 ng MEC-6 cRNA, 3.75 or 7.5 ng MEC-2 cRNA, 3.75 or 5 ng cRNA of POML-1, (no difference was seen for the different MEC-2 and POML-1 injections and data were pooled). For the remainder of the experiments in Figures 7 and S6, oocytes were injected with 3.75 ng Myc::MEC-4(d) cRNA, 1 ng MEC-6::HA cRNA, 10 ng MEC-2 cRNA, 7.5 ng cRNA of EGFP::POML-1 unless noted. Co-expression of Myc::MEC-4(d) and tagged MEC-6 and POML-1 produced amiloride-sensitive currents that were similar to those of the co-expressed untagged proteins at different time points [for MEC-4(d) and MEC-6 vs Myc::MEC-4(d) and MEC-6::HA at 2 days: −0.8 ± 0.1 vs −0.7 ± 0.1 and at 5-6 days: −1.7 ± 0.3 vs −1.7 ± 0.3; for MEC-4(d) and POML-1 vs Myc::MEC-4(d) and EGFP::POML-1 at 2 days −0.1 ± 0 vs −0.2 ± 0.1 and at 5-6 days: −0.7 ± 0.2 vs −0.8 ± 0.1. n=5 oocytes].

Immunoprecipitation was performed 5-6 days after cRNA injection as previously described (Goodman *et al*., 2002). Protein complex was precipitated by using the following antibodies conjugated to Protein A/G PLUS-Agarose (Santa Cruz Biotechnology, Dallas, Texas): antibodies against GFP (rabbit polyclonal sc-8334, Santa Cruz Biotechnology), Myc (rabbit polyclonal sc-789, Santa Cruz Biotechnology), and HA tags (rabbit polyclonal, sc-805, Santa Cruz Biotechnology). Protein samples were subjected to SDS-PAGE and Western blot. 4-8 oocyte equivalents were loaded per lane for immunoprecipitation, and one oocyte equivalent was loaded per lane for total lysate. The specificity of the immunoprecipitation was confirmed in two ways. First, 1 ng of cRNA encoding Myc::EGFP [for Myc::MEC-4(d) immunoprecipitation of EGFP::POML-1], EGFP [for EGFP::POML-1 immunoprecipitation of Myc::MEC-4(d)], and EGFP::HA (for MEC-6::HA immunoprecipitation of EGFP::POML-1) were used as negative controls; none of the proteins were immunoprecipitated. Second, we probed the immuno-complexes for a *Xenopus* oocyte membrane protein, β-integrin, by using a monoclonal antibody (8C8, Developmental Studies Hybridoma Bank, University of Iowa, IA), and did not detect the β-integrin.

Protein was detected by Western blot using antibodies against GFP (mouse monoclonal, sc-9996, Santa Cruz Biotechnology), Myc (mouse monoclonal 9E10, Sigma), the HA tags (mouse monoclonal, sc-7392, Santa Cruz Biotechnology), MEC-2 N-terminus (rabbit polyclonal) (Zhang, 2004), or actin (rabbit polyclonal, A2066, Sigma) and horse radish peroxidase (HRP)-conjugated secondary antibodies (Jackson ImmunoResearch Laboratories). Horseradish peroxidase was detected using the ECL Western Blotting reagent (Amersham).

Biotinylation assays to detect the surface expression of MEC-4 generally followed the methods described previously (Goodman *et al*., 2002). Surface protein was labeled and isolated using the membrane impermeable EZ-Link Sulfo-NHS-SS-Biotin and NeutrAvidin agarose provided in the Pierce Cell Surface Protein Isolation Kit (Thermo Scientific, Waltham, MA). Samples collected from 30 oocytes from each group were loaded per lane in SDS-PAGE and detected by Western blotting using a primary monoclonal Myc antibody (clone 9E10, Sigma) and a HRP-conjugated secondary antibody (Jackson ImmunoResearch Laboratories). The total lysate of one oocyte equivalent was loaded as input. β-actin was detected in total lysate as an input control by Western blotting using a rabbit polyclonal antibody against actin (A2066, Sigma). Cytoplasmic EGFP was detected in the supernatants but not in the avidin precipitates (not shown). Band density was measured from the autoradiography films using Image J.

TIRF imaging of oocytes was performed as described in Chen *et al*., 2015.

### Statistics

Data are presented as mean ± S.E.M unless noted. Statistical significance was determined using GraphPad Prism5 software (http://www.graphpad.com/scientific-software/prism/). We used the Student’s t test (with Welch’s correction when data being compared did not have equal variances) for most experiments. For the quantification of MEC-4::EGFP spots on the surface of *Xenopus* oocytes, we used the Mann-Whitney test. For the quantification of Western blot, we used the one sample t test. All p values from the Student’s t, Mann-Whitney, and one-sample t test were adjusted with a Bonferroni correction. When three or more groups were compared, we used oneway ANOVA with Tukey post hoc or two-way ANOVA with Bonferroni post-tests. In the figures *, **, and *** indicate corrected p values of <0.05, <0.01, and p<0.001, respectively; ns indicates not significant. All values were determined with the Student’s t test unless noted.

## SUPPLEMENTAL MATERIALS

Supplemental Materials include figures, a video and supplemental materials and methods can be found with this article online at …

## ACKNOWLEDGMENTS

We thank Jian Yang and Yong Yu for providing the *Xenopus laevis* oocytes, Mei Zhen for the puncta analysis software, and Oliver Hobert, Elizabeth Miller, and members of the Chalfie lab for discussion. This work was supported by grants GM30997 to M.C. and NS35549 to E.Y.I. from the National Institutes of Health.

